# Hobrac: a reference-guided workflow for genome comparison and synteny visualization

**DOI:** 10.64898/2026.07.17.739168

**Authors:** Benjamin Istace, France Denoeud, Emilie Téodori, Noor Chorba, Jean-Marc Aury

## Abstract

Whole-genome comparison is fundamental for validating genome assemblies and investigating genome evolution, yet identifying suitable reference genomes and interpreting chromosome-scale synteny from often noisy nucleotide alignments remain challenging. We introduce Hobrac, an automated workflow that addresses these two major bottlenecks by combining automated reference genome selection with gene-based structural comparisons. Starting from a genome assembly and its taxon identifier, Hobrac identifies suitable reference genomes, complements nucleotide alignments with conserved BUSCO orthologues, and generates publication-quality visualizations. The workflow produces dotplots, ribbon-plots and synteny visualization that can be explored interactively or offline. Hobrac is freely available at https://github.com/Genoscope-LBGB/hobrac.

## Background

Thanks to the advent of long-read and long-range sequencing, the number of reference genomes in public databases is increasing rapidly [1,2], notably with the help of large sequencing projects such as the Darwin Tree of Life [3], ATLASea [4] or the Vertebrate Genome Project [5]. Comparing these novel genome assemblies to established references is a common task in genomics that serves multiple purposes, including genome assembly validation, synteny analysis, chromosome rearrangement detection, and comparative studies of genome evolution. As the number of available reference genomes continues to grow, however, these analyses have become increasingly challenging. Public databases now contain hundreds to thousands of genomes for many taxonomic groups, making automated and reproducible workflows essential in place of manual reference selection and pairwise comparisons.

Current approaches rely on whole-genome alignments, typically represented as two-dimensional dotplots that highlight regions of similarity between genomes [6,7]. These genome-to-genome alignments are often noisy and difficult to analyze, especially when no closely related reference genome is available. Selecting an appropriate reference for structural comparison purposes is also often non-trivial, requiring manual exploration of the taxonomic neighborhood and iterative alignments to identify sufficiently close genomes. Although powerful alignment and visualization tools exist, they generally assume that suitable reference genomes have already been identified and provide limited support for automated comparative analyses across large genome collections.

Furthermore, less than 20% of chromosome-scale assemblies currently available in the NCBI Genbank database are annotated. Synteny and comparative genomics analyses generally rely on genome annotation and the identification of orthologous genes [8], which can substantially delay downstream analyses of assembled genomes. Conserved chromosome organization is also difficult to interpret. Whether projecting previously described ancestral linkage groups (ALGs) [9] onto new genome assemblies or inferring them *de novo* across diverse lineages, researchers often need to manually reconstruct chromosome correspondences because published ALG definitions are not directly transferable.

Here we present Hobrac (Homology-based Reference genome Acquisition and Comparison), a new tool designed to address these limitations in large-scale comparative genomic analyses. Hobrac automatically identifies suitable reference genomes from a curated database built from chromosome-level assemblies available in GenBank. This automated reference selection ensures that analyses remain reproducible while continuously benefiting from newly released reference genomes. It then uses the positions of conserved BUSCO genes [10,11] in the assembly and the reference to generate dotplot visualizations. These conserved genes provide robust anchors for structural comparisons across larger evolutionary distances, reducing the analysis to a few thousand loci instead of millions of nucleotide alignments and greatly simplifies the interpretation of chromosome-scale relationships. Whole-genome alignments are also generated to complement gene-based comparisons whenever nucleotide-level similarity remains informative.

Besides dotplots, Hobrac generates ribbon plots that show synteny blocks across multiple genomes simultaneously. These plots highlight ALGs, i.e. chromosomal units that have retained conserved gene linkages over evolutionary time [9,12–15]. ALGs can either be inferred directly from the compared genomes or projected from previously described ancestral karyotypes. In addition, HoBRAC computes a rearrangement index that quantifies the extent of chromosome rearrangements between each genome and the reference ALGs, providing a complementary measure of structural conservation.

Finally, we developed a web-based alignment viewer that accepts any PAF alignment file, allowing users to explore genome comparisons generated either by Hobrac or by external alignment workflows.

We illustrate the versatility of Hobrac through a series of genomics applications. These include assembly validation using conserved gene-based structural comparisons, as well as chromosome-scale evolutionary analyses based on multi-genome synteny visualization, *de novo* reconstruction of ALGs, and projection of previously defined ALGs onto newly assembled genomes.

## Results

### Hobrac overview

Hobrac is an automated comparative genomics workflow that requires only a genome assembly and its taxon identifier to perform a series of analyses, from automated reference selection to chromosome-scale synteny visualization, supporting both genome assembly validation and comparative genomics (Figure 1). One or several reference genomes are first selected using Mash, an alignment-free method that summarizes each genome’s k-mer content as a compact MinHash sketch. The overlap between sketches is then used to rapidly estimate genomic distance, allowing the input assembly to be screened efficiently against a phylum-specific database that we built from chromosome-level Genbank assemblies. The closest genomes are then downloaded from NCBI and complete single-copy BUSCO genes are identified in the query assembly and the selected references to serve as synteny markers. In parallel, Minimap2 [16] whole-genome alignments provide nucleotide-level comparisons and a higher-resolution view whenever sequence similarity is sufficient.

**Figure 1.**
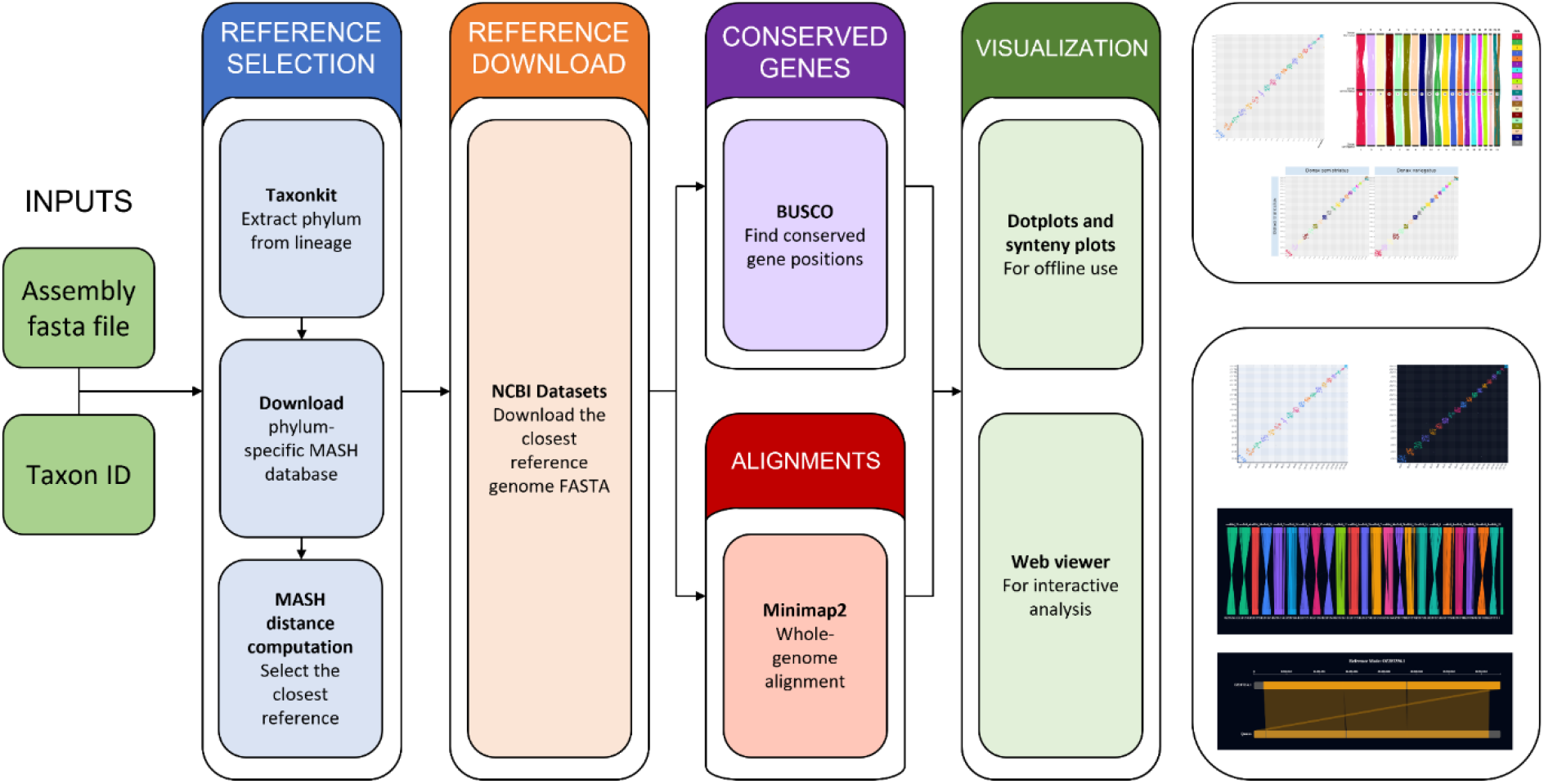
Description of the Hobrac workflow. Hobrac takes a genome assembly and a taxon identifier as inputs, then proceeds through five stages: reference genome selection using Mash distance against phylum-specific databases, reference genome download via NCBI datasets, conserved gene identification using BUSCO on both query and reference, whole-genome alignment with Minimap2, and visualization through static dotplots and synteny plots or an interactive web interface.

The resulting gene- and nucleotide-level alignments support two complementary applications. For assembly inspection and validation, BUSCO-based dotplots provide a simplified representation of chromosome correspondences that is easier to interpret than whole-genome alignments. For comparative genomics, HoBRAC provides two complementary analyses: ancestral linkage group (ALG) inference and rearrangement index computation. Hobrac generates ribbon plots and synteny grids that display conserved BUSCO genes colored according to their ALG. These ALGs can either be inferred (i) *de novo* by testing if shared BUSCO genes co-occur more often than expected with a Fisher exact test and by grouping significant associations into chromosome chains, or (ii) projected from previously described datasets using precomputed BUSCO colour schemes covering for instance the previously described/known 29 metazoan and 24 bilaterian ALGs [9]. Hobrac also allows users to provide their own BUSCO-to-colour files to visualize alternative or custom chromosome classifications.

Hobrac is written in Python and Snakemake [17] and is available at https://github.com/Genoscope-LBGB/hobrac. The visualizer has been built with React and the d3.js library and is available at https://www.genoscope.cns.fr/lbgb/hobrac. At the time of writing, the Hobrac reference genome database is composed of 16,083 chromosome-level assemblies available in NCBI’s Genbank database, spanning 40 phyla. The number of reference genomes available in each phylum varies greatly, with Streptophyta (4,362 genomes), Chordata (4,221 genomes) and Arthropoda (3,894 genomes) being the most represented while some others like Bryozoa (8 genomes) and Ctenophora (5 genomes) having less than 10 reference genomes. For each phylum, we computed a Mash sketch database that makes it possible to rapidly compute the distance of any genome assembly to all the reference genomes present in the database. Additionally, mappings of already described ALGs on BUSCO genes are available for 9 BUSCO odb12 datasets: actinopterygii, vertebrata, lophotrochozoa, mollusca, crustacea, anthozoa, cnidaria, metazoa and arthropoda.

### Using Hobrac for assembly validation

To illustrate Hobrac for assembly validation, we analyzed the chromosome-scale assembly of *Donax trunculus*. Hobrac automatically identified the closest reference genome available in NCBI’s Genbank: *Donax semistriatus* (Mash distance of 0.1744). On our Slurm cluster the complete analyses, from reference genome selection to the generation of dotplots and ribbon plots, completed in 34 minutes (8.46 CPU hours) with a peak memory usage of 23.9GB corresponding to the peak memory usage of Minimap2. Most of the computational time was spent running BUSCO (7.91 CPU hours), whereas whole-genome alignment with Minimap2 required only 0.51 CPU hours.

Using the Mollusca BUSCO dataset, Hobrac identified 4,250 single-copy orthologues shared between the two assemblies. The resulting BUSCO-based dotplot clearly resolved one-to-one chromosome correspondences, with all 20 syntenic chromosome pairs immediately identifiable. In contrast, the corresponding whole-genome alignment produced a substantially noisier dotplot despite showing the same overall chromosome organization (Figure 2).

**Figure 2.**
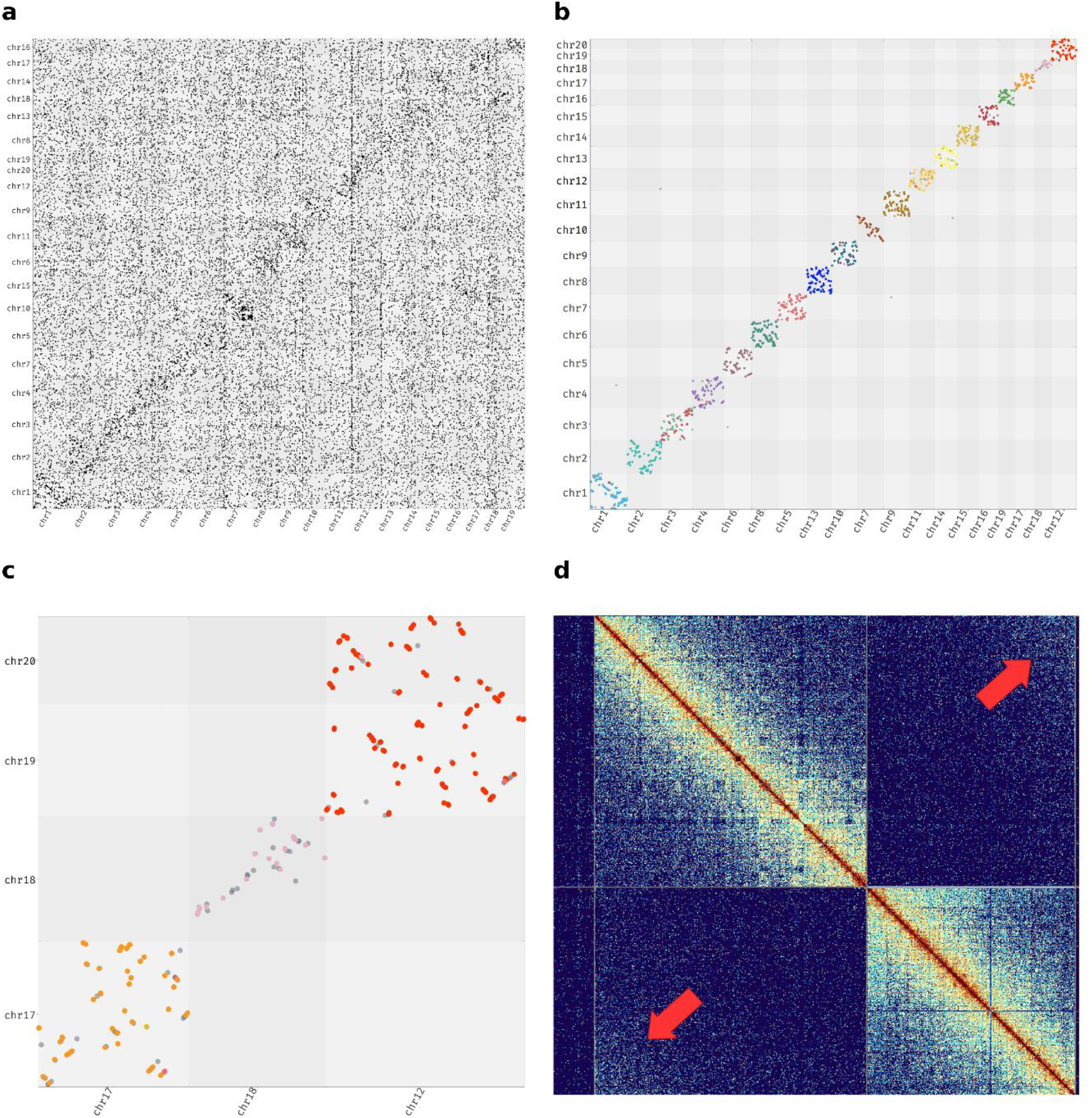
Use of Hobrac for assembly curation. **a**. Whole-genome alignment dotplot of *Donax trunculus* (y-axis) against *Donax semistriatus* (x-axis). **b**. BUSCO-based dotplot of *Donax trunculus* (y-axis) against *Donax semistriatus* (x-axis), points are colored based on the known 24 bilaterian ALGs. **c**. Zoomed view of the BUSCO-based dotplot suggesting the possible fusion of chromosomes 19 of 20 of *Donax trunculus* (x-axis) based on chromosome 12 of *Donax Semistriatus*. **d**. Hi-C contact map of scaffolds 19 and 20 from *Donax trunculus*. Red arrows indicate regions of increased inter-chromosomal contacts, supporting the inversion of scaffold 20 and its subsequent joining with scaffold 19.

Interestingly, *D. semistriatus* contains 19 chromosome-scale scaffolds, whereas *D. trunculus* contains 20. Pairwise comparisons showed that both scaffolds 19 and 20 of *D. trunculus* have strong links with chromosome 12 of *D. semistriatus*. All remaining chromosomes showed one-to-one organization across both species.

The synteny analysis raised the possibility that scaffolds 19 and 20 of *D. trunculus* belonged to the same chromosome, leading us to inspect the Hi-C contact map (Figure 2d). Scaffolds 19 and 20 displayed inter-scaffold contacts consistent with them belonging to a single chromosome. Merging the two scaffolds restored the expected karyotype of 19 chromosomes previously described for this species [18]. This example demonstrates how conserved gene-based structural comparisons can effectively identify candidate assembly errors that may remain difficult to recognize using nucleotide-level alignments alone and can subsequently be validated with orthogonal data such as Hi-C.

### Using Hobrac for synteny analyses

To illustrate the use of Hobrac for comparative synteny analyses, we ran the workflow with selected references on seven bivalve species of the subclass Autobranchia. The dataset contains six Heteroconchia: *Donax trunculus, Donax semistriatus, Gari depressa, Laevicardium crassum, Tridacna crocea*, and *Venus verrucosa*, and one Pteriomorphia: *Pecten maximus*. On our Slurm cluster the entire process completed in 31 minutes (20.16 CPU hours) with a peak memory usage of 20.9GB corresponding to the peak memory usage of BUSCO. The running time was mainly explained by the different BUSCO steps which completed in a total of 20.13 CPU hours.

On the seven bivalve genomes, Hobrac built 20 ALGs *de novo*. The six Heteroconchia genomes share a fusion of two ALGs (“18” and “19”), whereas the Pteriomorphia *Pecten maximus* underwent a separate fusion that independently led to 19 chromosomes. The ribbon plot shows that the chromosome structure is well conserved across Heteroconchia, with only one additional fusion event in *Tridacna crocea* (Figure 3a). These ribbon plots provide a concise overview of chromosome conservation and lineage-specific rearrangements across multiple genomes.

**Figure 3.**
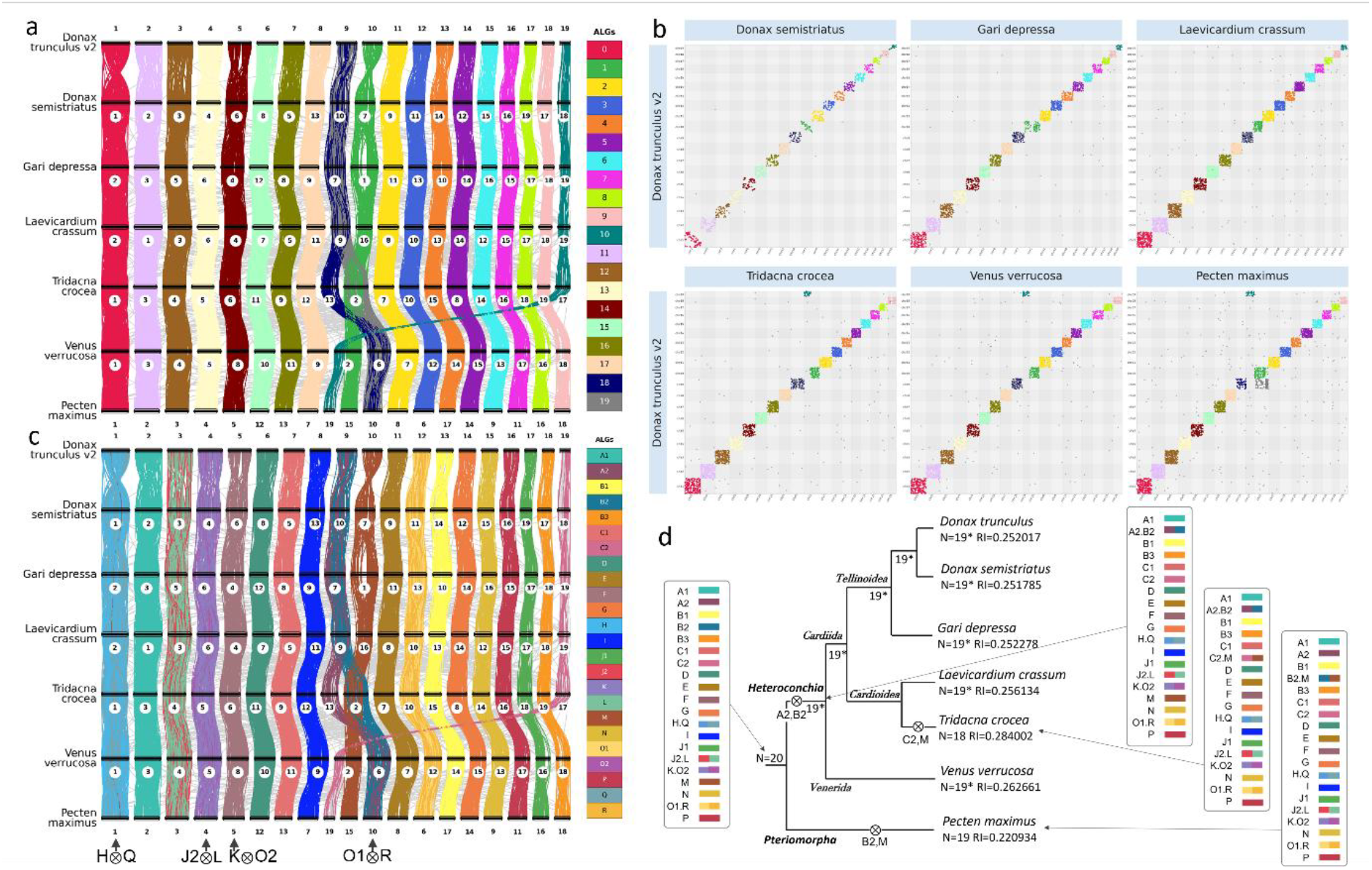
Use of Hobrac for synteny analyses. a. Synteny ribbon plot obtained for 7 bivalve genomes. Colors correspond to the ALGs identified by Hobrac with a significance threshold of P=0.05. b. Dotplot grid corresponding to the ribbon plot in a, using the same coloring scheme. c. Ribbon plot generated using the option to color the genes according to the previously described bilaterian ALGs. Arrows below the plot indicate the fusion-with-mixing events leading from the 24 bilaterian ALGs to the 20 molluscan ALGs. d. Evolutionary scenario inferred from the synteny plots, illustrating the transitions from the 24 bilaterian ALGs to the extant karyotypes of the seven bivalve species. The tree represents a schematic phylogenetic topology (branch lengths are not to scale). Fusion-with-mixing events are indicated by ⊗. Species sharing the same chromosome structure are noted with *.

An alternative way of viewing synteny is through pairwise comparison of genomes with dotplots. Hobrac builds grids of dotplots from the BUSCO gene positions colored by ALG, using the same colour scheme as in the ribbon plots. On those plots, the “assembly” genome is put on the y-axis as a reference for all comparisons. This complementary representation facilitates the identification of chromosome fusions, fissions, whole-genome duplications and fusion-with-mixing events (noted with ⊗ as initially proposed by Simakov et al [9]). Dotplots complement ribbon plots by making local gene order more readily apparent, thereby facilitating the simultaneous analysis of macro- and microsynteny. For example, conserved gene order remains clearly visible between *D. trunculus* and *D. semistriatus* chromosome pairs 10-7 and 19-18 (Figure 3b).

While *de novo* inference of ALGs provides a simplified view of chromosomal architecture, Hobrac also supports the projection of previously described ALGs onto newly analysed genomes, enabling direct evolutionary interpretation in the context of published ancestral karyotypes. To achieve this, we computed BUSCO-to-ALG colour mappings for several BUSCO datasets, allowing ribbon plots and dotplots to be coloured according to known ALGs [9,19]. The synteny plots generated on the seven bivalve genomes with the option dedicated to colouring the 24 bilaterian ALGs revealed four fusion-with-mixing events shared by all analysed species: H⊗Q, K⊗O2, J2⊗L and O1⊗R (Figure 3c). These correspond to the four ancestral chromosome fusions previously described as shared by Lophotrochozoa and possibly already present in the spiralian ancestor [9,20], which gave rise to the 20 ALGs of the molluscan ancestor [21]. The 20 ALGs identified *de novo* by Hobrac (Figure 3a) exactly correspond to these molluscan ALGs, demonstrating that the *de novo* approach independently recovers the previously inferred ancestral molluscan karyotype. Finally, the synteny plots facilitate the reconstruction of the history of chromosomal rearrangements leading from the 20 molluscan ALGs to the karyotypes of the seven extant species (Figure 3d). Three fusion-with-mixing events occurred: B2⊗M in the lineage leading to *Pecten maximus*, A2⊗B2 in the ancestor of all Heteroconchia and C1⊗M in the lineage leading to *Tridacna crocea*. The computation of rearrangement indexes (RI) with respect to the provided ALGs was implemented in Hobrac, with the procedure proposed by Lewin et al. [19,22]. As expected, *Tridacna crocea*, which encountered an additional fusion compared to other Heteroconchia, displays the highest RI (0.284) among the species compared. *Pecten maximus*, on the other hand, has the lowest RI (0.221), confirming its suitability as a reference mollusc genome for synteny analyses.

## Discussion

Hobrac was developed to facilitate structural comparison of newly assembled genomes with existing references, without requiring prior gene annotation. By relying on conserved single-copy BUSCO genes, it can be applied immediately after assembly and provides a simplified representation of chromosome correspondences that is often easier to interpret than whole-genome alignments alone. In our experience, this makes it particularly useful as a support for assembly validation and manual curation, for example during the interpretation of Hi-C contact maps or for identifying sex chromosomes. Beyond the individual methods it combines, the main contribution of Hobrac lies in integrating automated reference selection, gene- and nucleotide-level comparisons, ALG inference and interactive visualization into a single reproducible workflow, thereby substantially reducing the manual effort traditionally required for comparative genomics.

Although Hobrac is primarily intended for chromosome-scale assemblies, it can also provide useful information for more fragmented genomes. In such cases, conserved gene correspondences may help identify scaffolds likely belonging to the same chromosome and generate hypotheses for manual scaffolding when long-range information is not available. However, these hypotheses should be interpreted with caution, as deviations from the reference genome may reflect genuine lineage-specific chromosomal rearrangements rather than assembly errors and should therefore be validated using orthogonal data. The interpretability of the results strongly depends on assembly contiguity, as fragmented assemblies reduce the continuity of the synteny signal and complicate structural inference.

Another advantage of Hobrac is that although previously described ALGs provide a useful framework for evolutionary interpretation, it does not depend on their prior existence. In lineages where deep ancestral chromosomal organization has been extensively reshuffled, synteny conservation can still be detected between closely related species. In such cases, the *de novo* inference of ALGs provides a simplified representation of recurrent chromosome associations and may help identify lineage-specific structural conservation. Because *de novo* inferred ALGs are defined from the set of genomes included in a given analysis, their composition varies across datasets. To facilitate their reuse, HoBRAC outputs can be used to generate custom BUSCO-to-colour files that preserve ALG assignments across subsequent analyses.

We discuss below several limitations of Hobrac and potential avenues for improvement. First, Hobrac automates reference selection for structural comparison between genomes using Mash distances, but this relies on the assumption that genomic similarity reflects structural conservation, which is not necessarily the case. Indeed, closely related genomes may differ substantially in chromosome organization because of lineage-specific rearrangements, while more distant genomes may sometimes retain more informative synteny patterns. This also implies that the closest genomes according to Mash are not always the closest in evolutionary terms and several reference genomes can have nearly identical Mash distances while being structurally distant. Future versions of Hobrac could integrate additional criteria for reference ranking, such as assembly quality, chromosome completeness or biological metadata including sex, thereby improving comparisons involving heteromorphic sex chromosomes.

Hobrac relies exclusively on complete single-copy orthologs, so the resolution of the analysis is constrained by the number and phylogenetic relevance of the available BUSCO markers. In poorly represented phyla, BUSCO datasets may contain fewer informative genes reducing the signal available for structural comparison. However, our ability to map known ALGs on various BUSCO datasets using phylogenetically diverse metazoan genomes demonstrates that even distant BUSCO datasets can remain informative enough to recover meaningful chromosome correspondences. Moreover, although the choice of using complete single-copy BUSCO genes simplifies orthology relationships and improves the interpretability of the comparisons, this reduces the applicability of the method to polyploid genomes, assemblies containing unresolved haplotypes, or lineages that underwent recent whole-genome duplications. Extending the framework to incorporate duplicated BUSCO genes could broaden its applicability to these cases and provide a more informative representation of genome structure in duplicated genomes.

The BUSCO-based representation also imposes intrinsic resolution limits. Because the approach focuses on conserved coding regions, rapidly evolving, gene-poor, or repeat-rich genomic regions are underrepresented. Structural changes occurring in these regions may therefore remain undetected. For this reason, Hobrac complements gene-based synteny analyses with whole-genome nucleotide alignments, which provide higher-resolution views of genome structure when sequence similarity remains sufficient. Rather than replacing whole-genome alignments, Hobrac provides an additional level of abstraction that simplifies chromosome-scale interpretation while retaining access to nucleotide-level comparisons when finer structural resolution is required. The two approaches are therefore best considered complementary rather than interchangeable.

## Conclusions

Hobrac provides a reproducible framework that integrates automated reference selection, conserved gene- and nucleotide-level structural comparisons, ancestral linkage groups analyses and interactive visualization. By complementing whole-genome alignments with chromosome-scale representations based on conserved BUSCO orthologues, Hobrac facilitates genome assembly curation, comparative synteny analyses and the reconstruction of chromosome evolution across diverse lineages.

As chromosome-scale genome assemblies continue to accumulate through large-scale sequencing initiatives, automated and reproducible comparative genomics workflows will become increasingly important. As Hobrac requires only a genome assembly and its taxonomic identifier it provides a practical solution to process large collections of assemblies with minimal manual intervention. Its multi-reference mode further helps distinguish lineage-specific chromosomal rearrangements from potential assembly artefacts, while the standalone visualization interface can also be used independently with any PAF alignment generated by external workflows.

## Methods

### Reference genomes database construction

In order to compare a genome assembly to publicly available genomes, we first downloaded all chromosome- or complete-level assemblies from the NCBI Genbank database and partitioned these genome assemblies by their phylum. We then created a Mash database for each phylum, based on the input genome assemblies. The phylum extraction was done with TaxonKit [23], and Mash v2.3 [24] was run with a sketch size of 10,000 (-s 10000).

### Automated reference selection

When a genome assembly is given as an input to Hobrac, its entire lineage is extracted using TaxonKit and the taxon identifier that the user provides. The Mash database that corresponds to the genome assembly’s phylum is then downloaded and the Mash distance between the input genome assembly and all other reference genomes is computed. The genome assembly with the lowest Mash distance, representing the highest genomic similarity, to the assembly is selected as the closest reference genome. Multiple references can be selected when specified by the user, and users may alternatively provide custom reference genomes directly in fasta format. Hobrac also contains options to avoid selecting species with the same taxon identifier or genomes with a Mash distance of zero, as such nearly identical genomes are unlikely to provide informative structural comparisons.

### Conserved genes identification

Hobrac automatically selects the most appropriate BUSCO dataset by traversing the taxonomic hierarchy from species to kingdom and matching against available OrthoDB lineages. BUSCO is then run in genomic mode on both the genome assembly and the selected reference. Genomic coordinates of complete single-copy orthologs are then extracted and converted to PAF (Pairwise Alignment Format), where each ortholog pair is represented as an alignment anchored at the gene midpoint to enable synteny visualization.

### Ancestral Linkage Group (ALG) detection

For each pair of species, the number of shared single-copy BUSCO genes is counted on every chromosome pair. A Fisher exact test with Bonferroni correction is then applied to determine whether the observed co-occurrence of genes on a given pair of chromosomes is significant, and only associations with a p-value below 0.01 by default are retained.

To ensure that ALG colors remain consistent when comparing the structure of multiple species, significant associations are organized into chromosome chains. Each BUSCO gene is walked through the species along its successive chromosomes, skipping species where the gene is absent. Maximal paths in which all edges correspond to significant associations define candidate chains. Chains supported by fewer than five genes are discarded by default.

The remaining chains are then subjected to a validation step. Indeed, each node in a chain of length n must have significant associations with at least ⌊n/2⌋ other nodes within the same chain. Nodes that do not satisfy this criterion are pruned, and contiguous segments spanning at least two species are retained as final ALGs. Each ALG receives a color that is maintained across all pairwise comparisons. A more permissive validation mode is also available, in which only one significant link per node is required regardless of chain length. Detected ALGs as well as gene chains and the ALG they are assigned to are written to a file so that users can inspect them. Alternatively, users can use the skip-alg option and provide a custom color file containing ALG assignments for each BUSCO gene in order to skip the detection step entirely.

### Mapping known ALG on BUSCO datasets

Color files corresponding to ALGs defined by Simakov et al. [9] were generated for various BUSCO odb12 datasets using the following procedure. We first ran Hobrac on genomes previously used for ALG identification [9,22] in order to identify the chromosome associations corresponding to the 29 “BCS” (bilaterian, cnidarian, sponges) ALG, that we will name “metazoan” ALGs for more simplicity, and the 24 bilaterian ALGs described by Simakov et al. [9], and derive rules to assign unambiguously the BUSCO genes to one of the 29 metazoan ALGs and the 24 bilaterian ALGs, from their chromosomal locations (Figure 4).

**Figure 4.**
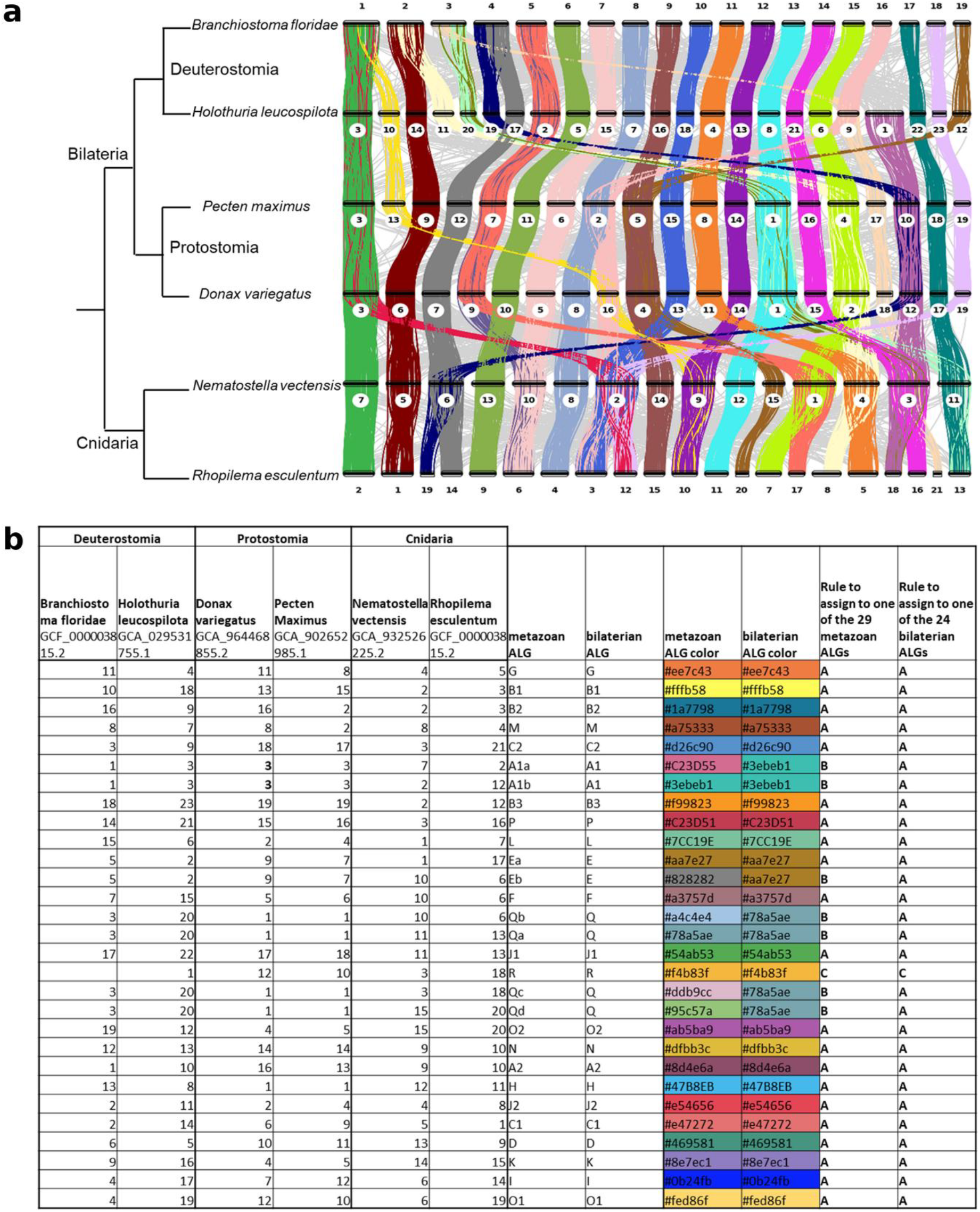
Mapping of previously described ALGs on BUSCO datasets. **a**. Ribbon plot obtained with Hobrac on six metazoan genomes using the BUSCO mollusca odb12 gene set. The tree represents a schematic phylogenetic topology (branch lengths are not to scale). ALGs were computed *de novo* with a significance threshold of P=0.05 for chromosome associations and the – permissive-alg option. **b**. Chromosomal associations corresponding to the 29 metazoan and 24 bilaterian previously described ALGs, and derived rules used to color BUSCO genes based on their chromosome location. Rules are the following: A. BUSCO gene found in at least three of the six genomes, and all chromosomes correspond to the ALG (or all but one if BUSCO gene found in >=5 genomes). B. BUSCO gene found in at least three of the six genomes, including at least one cnidaria, and all chromosomes correspond to the ALG (or all but one if BUSCO gene found in >=5 genomes). C. BUSCO gene found in at least three genomes, and all chromosomes correspond to the ALG, not considering Branchiostoma floridae(or all but one if BUSCO gene found in >=4 genomes) -> group “R”.

Then, we performed BUSCO searches using various BUSCO gene sets on the 6 genome assemblies and followed the rules defined to assign the BUSCO genes to metazoan and bilaterian ALGs. We used the colors from [25] to color these ALGs. Color files are available for nine BUSCO odb12 datasets, and available at https://github.com/Genoscope-LBGB/hobrac/tree/master/hobrac/colors.

### Rearrangement index computation

When the user provides ALGs via a custom color file, or chooses to use a pre-computed color file with the option “--color-metazoan-alg” or “--color-bilaterian-alg”, Hobrac computes the rearrangement index relative to the provided ALGs for each reference genome and the assembly, following the methodology established by Lewin et al. [19,22]. For each ALG defined in the color file, the index combines the proportion of ALG genes retained on the best-matching chromosome and the proportion of genes on that chromosome belonging to the ALG. Species-level values are calculated as the mean across ALGs present in the species.

### Whole-genome alignment

Hobrac also generates the genome-to-genome alignments of the provided genome assembly to the selected references using Minimap2 [16] with the asm20 preset. These alignments offer a more precise view of the structural differences between two genomes but are typically noisier than the conserved genes alignments, due to repetitive elements and particularly when comparing evolutionary distant genomes.

### Visualization

We provide two ways to easily analyze Hobrac results. Dotplots and ribbon plots are always computed for offline analysis when running Hobrac. To facilitate genome assembly curation, we have built a custom alignment visualizer for interactive visualization.

#### Static dotplot generation

For offline analysis, we built dotplotrs, a command-line tool written in Rust. Dotplotrs reads PAF-formatted files and outputs dotplots in PNG format. To facilitate interpretation, genome assembly sequences are ordered against reference genome chromosomes using two strategies. The default ordering employs a gravity-based algorithm adapted from DGenies [7], which ranks sequences by their alignment weight per target chromosome. Alternatively, a significance-based ordering employs a Fisher’s exact test to identify statistically enriched associations between queries and target sequences. P-values are corrected for multiple testing by using the Bonferroni method and associations with p-values inferior to 0.01 are considered significant. Significantly associated alignments with chromosomes are displayed in distinct colors while non-significant alignments appear in gray. Additional customization options include minimum alignment length filtering, line thickness adjustment, light or dark color schemes, custom alignment colors and custom sequence ordering. Dotplotrs is freely available on Github: https://github.com/Genoscope-LBGB/dotplotrs.

#### Ribbon plot visualization

Ribbon plots are generated using the karyotype module of the JCVI toolkit [8]. Chromosomes are displayed as horizontal tracks ordered by size, with synteny ribbons connecting orthologous BUSCO genes between adjacent species. Ribbons are colored according to ALG membership, with non-significant associations shown in grey. This visualization facilitates rapid identification of conserved chromosomal units, chromosome fusions, and fissions across the compared genomes. The resulting image is then post-processed to add a legend that the color of each ALG to their respective color.

#### Dotplot synteny grid

To complement the ribbon plots with a per-reference view, Hobrac assembles the individual BUSCO dotplots into a synteny grid. Each cell of the grid is a dotplot of the BUSCO genes shared between the assembly and one reference genome. Points are colored according to their ALG membership, using the same color scheme as the ribbon plots, while non-significant associations are shown in grey. Cells are tiled into a square-shaped grid and rendered at native resolution, which makes it possible to inspect every reference at once while remaining zoomable for closer examination.

#### Interactive web visualizer

For interactive exploratory analysis, we developed a web-based visualizer using React and d3.js. The application processes Hobrac whole genome alignment, BUSCO genes comparison or user-provided PAF files locally, without server-side data transmission, and builds three different views. The primary dotplot view shows sequences ordered by gravity and alignments colored either by identity or statistical significance. Users can pan, zoom, select specific sequences, or filter alignments in this view. The secondary ribbon view displays the assembly sequences above the reference sequences, with ribbons connecting aligned regions/genes according to the filters applied in the dotplot view. Finally, a third detail view can be activated by clicking individual alignments in the dotplot view, and shows all query sequences aligning to a selected reference chromosome. The visualizer is freely available at: https://www.genoscope.cns.fr/lbgb/hobrac/.

### Assembly validation

In order to find references for the *Donax trunculus* V1 genome (GCA_964273945.1), Hobrac was run in automatic mode and asked to find the closest reference genome by leaving the “--ref-count 1” option as default. The genome of *Donax semistriatus* (GCA_965239255.1) was the closest reference available in NCBI’s GenBank database according to Mash distance.

### Synteny analysis

The six reference genomes used for the bivalves synteny analyses were given manually to Hobrac via the use of the “--reference” option which disables the automatic Mash-distance-based search. The reference genomes that were used were *Donax semistriatus* (GCA_965239255.1), *Gari depressa* (GCA_965649845.1), *Laevicardium crassum* (GCA_964662305.1), *Tridacna crocea* (GCA_943736015.1), *Venus verrucosa* (GCA_964200665.2) and *Pecten maximus* (GCA_902652985.1) while the genome given as an ‘assembly’ was *Donax trunculus* V2 (GCA_964273945.3). The Minimap2 alignments were skipped as the focus was on synteny by using the “--skip-genomic” option. The Fisher’s test p-value was relaxed to 0.05 with “--alg-pvalue 0.05” and sequences needed at least 30 BUSCO genes to be considered for ALG construction thanks to the “--min-busco-genes 30” option.

A second run was performed with the same input genomes and the additional option “–color-bilaterian-alg” in order to color the BUSCO genes according to the previously described bilaterian ALG they were assigned to. The option –skip-alg was not used, which means that the only links between genes that are detected as part of ALGs by the ab initio approach are colored in the ribbon plot (others are light grey).

## Availability of data and materials

Hobrac is an open source software, source code and binaries are freely available from https://github.com/Genoscope-LBGB/hobrac.

## Competing interests

The authors declare that they have no competing interests.

## Funding

This work was supported by the Genoscope, the Commissariat à l’Énergie Atomique et aux Énergies Alternatives (CEA), France Génomique (ANR-10-INBS-09-08), and the exploratory research programme ‘ATLASea: Atlas of marine genomes and its targeted projects DIVE-Sea (ANR-22-EXAT-0002-DIVE-Sea), SEQ-Sea (ANR-22-EXAT-0003-SEQ-Sea) and BYTE-Sea (ANR-22-EXAT-0004-BYTE-Sea).

## Authors’ contributions

JMA and BI proposed the initial concept to use both nucleotide- and protein-level alignments. BI conceived and implemented the Hobrac pipeline. FD mapped the existing ALGs on BUSCO datasets and conceived the synteny analysis part of the pipeline. BI and FD conducted the experiments. JMA supervised the analyses. BI, FD and JMA authored and reviewed drafts of the article. ET tested the pipeline and the online visualizer. NC implemented the draft version of the offline dotplot software. All authors read and approved the final manuscript.

## Notes

### Competing Interest Statement

The authors have declared no competing interest.

### Summary of Updates

Fixed a typo error in the biorxiv abstract preview (BUSCO rthologues -> BUSCO orthologues)

